## Supplementary materials for "High-level cognition during story listening is reflected in high-order dynamic correlations in neural activity patterns"

June 10, 2021

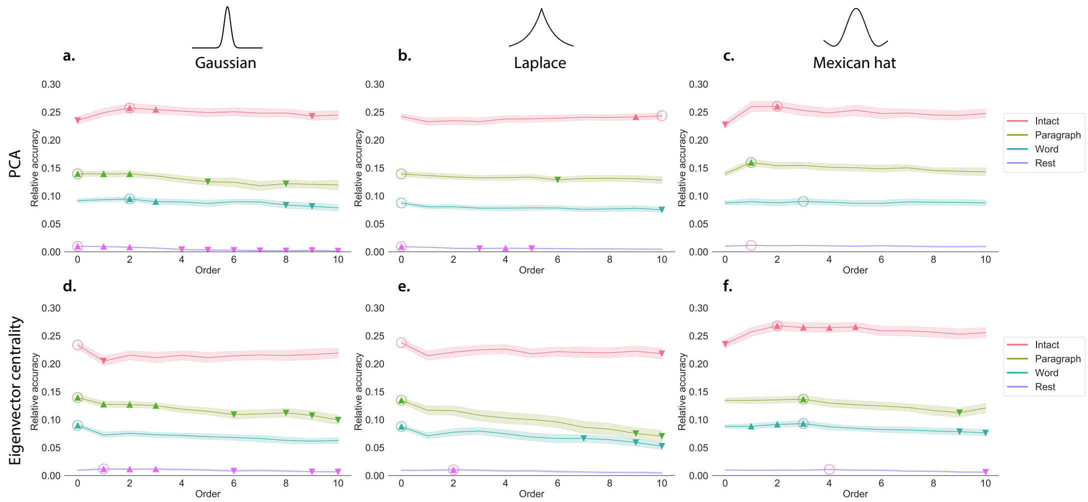


**Figure S1: Across-participant timepoint decoding accuracy varies with correlation order and cognitive engagement across kernels. a.-c. Timepoint decoding accuracy as a function of order: PCA.** *Order* ($x$-axis) refers to the maximum order of dynamic correlations that were available to the classifiers (see *Feature weighting and testing*). The reported across-participant decoding accuracies for **a. Gaussian**, **b. Laplace**, and **c. Mexican hat** kernel shapes are averaged over all widths (see *Identifying robust decoding results*). The $y$-values are displayed relative to chance accuracy (intact: $\frac{1}{300}$; paragraph: $\frac{1}{272}$; word: $\frac{1}{300}$; rest: $\frac{1}{400}$). The error ribbons denote 95% confidence intervals across cross-validation folds (i.e., random assignments of participants to the training and test sets). The colors denote the experimental condition. Arrows denote sets of features that yielded reliably higher (upward facing) or lower (downward facing) decoding accuracy than the mean of all other features (via a two-tailed $t$-test, thresholded at $p < 0.05$). The circled values represent the maximum decoding accuracy within each experimental condition. Panels a.-c. used PCA to project each high-dimensional pattern of dynamic correlations onto a lower-dimensional space. **d.-f. Timepoint decoding accuracy as a function of order: eigenvector centrality.** These panels are in the same format as Panel a.-c., but here eigenvector centrality has been used to project the high-dimensional patterns of dynamic correlations into lower-dimensional spaces.

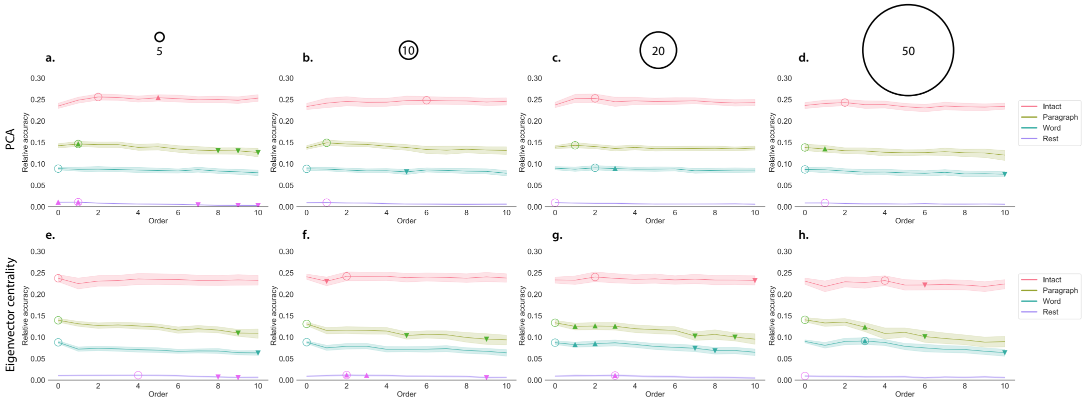


**Figure S2: Across-participant timepoint decoding accuracy varies with correlation order and cognitive engagement across widths. a.-d. Timepoint decoding accuracy as a function of order: PCA.** *Order* ($x$-axis) refers to the maximum order of dynamic correlations that were available to the classifiers (see *Feature weighting and testing*). The reported across-participant decoding accuracies for kernel widths of **a. 5**, **b. 10**, **c. 20**, and **d. 50** are averaged over all kernel shapes (see *Identifying robust decoding results*). The $y$-values are displayed relative to chance accuracy (intact: $\frac{1}{300}$; paragraph: $\frac{1}{272}$; word: $\frac{1}{300}$; rest: $\frac{1}{400}$). The error ribbons denote 95% confidence intervals across cross-validation folds (i.e., random assignments of participants to the training and test sets). The colors denote the experimental condition. Arrows denote sets of features that yielded reliably higher (upward facing) or lower (downward facing) decoding accuracy than the mean of all other features (via a two-tailed $t$-test, thresholded at $p < 0.05$). The circled values represent the maximum decoding accuracy within each experimental condition. Panels a.-d. used PCA to project each high-dimensional pattern of dynamic correlations onto a lower-dimensional space.**e.-h. Timepoint decoding accuracy as a function of order: eigenvector centrality.** These panels are in the same format as Panel a.-d., but here eigenvector centrality has been used to project the high-dimensional patterns of dynamic correlations into lower-dimensional spaces.

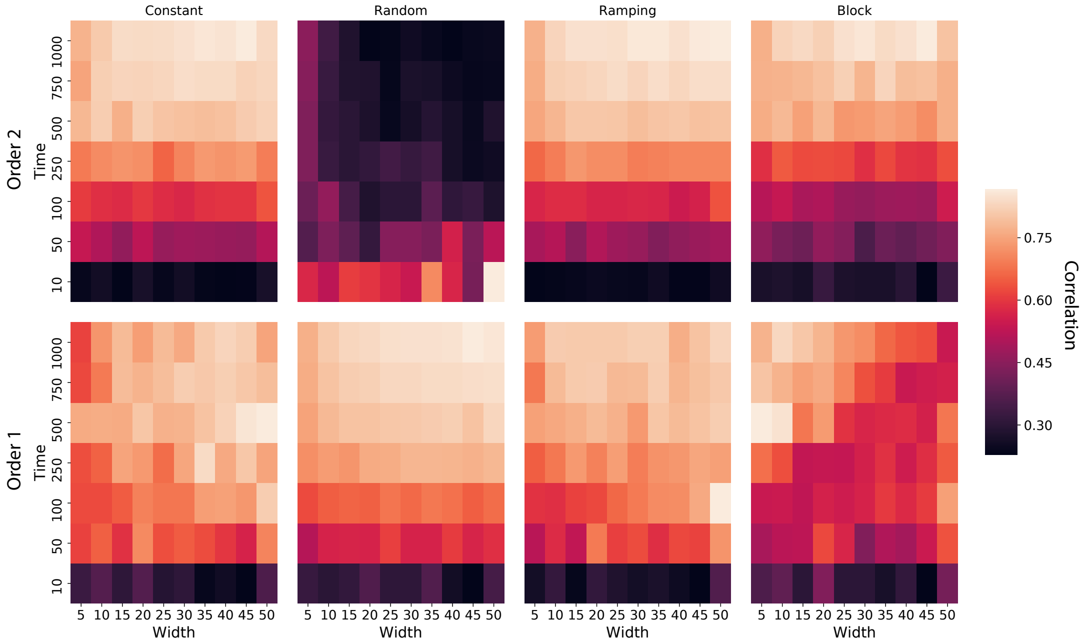


**Figure S3: Recovery heatmap of simulated first-order and second-order dynamic correlations across time and widths.** Each panel displays a heatmap of the average correlations between the vectorized upper triangles of the recovered first-order and second-order correlation matrices and the true (simulated) first-order and second order correlation matrices. The averages are taken across several randomly generated synthetic datasets for each timeseries pattern (constant, random, ramping, and data; see *Synthetic data: simulating dynamic higher-order correlations*). The $x$-axes of each heatmap denote varying Laplace-shaped kernel widths (see Figs. S1 and S2), and the $y$-axes of each heatmap denote varying durations (in samples) of the synthetic datasets. A total of 10 synthetic datasets were generated for each duration (row) and timeseries pattern (panel).

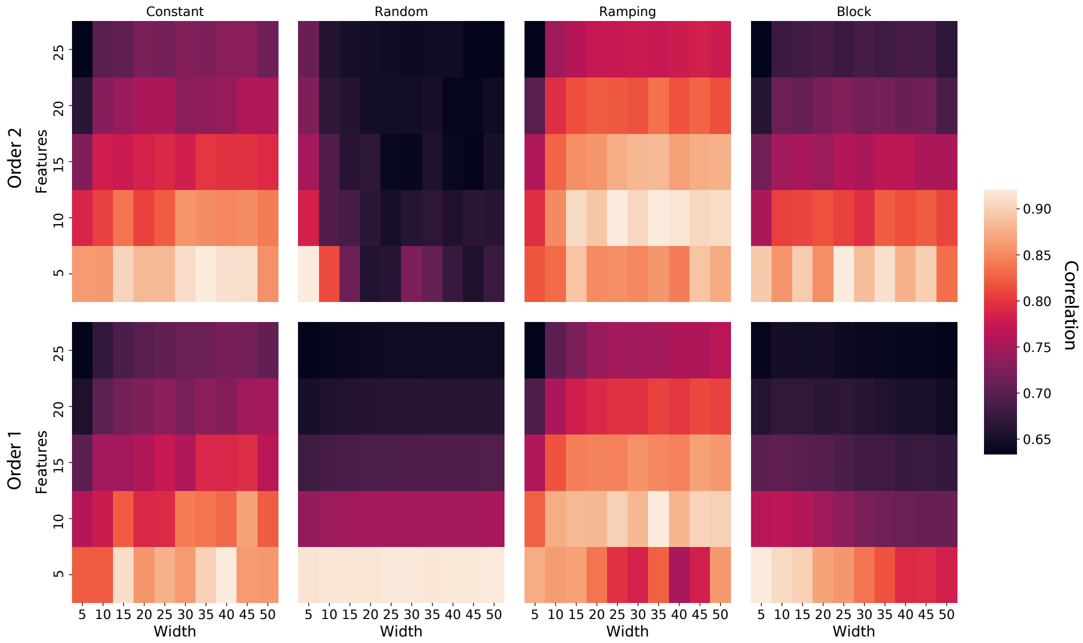


**Figure S4: Recovery heatmap of simulated first-order and second-order dynamic correlations across features and widths.** Each panel displays a heatmap of the average correlations between the vectorized upper triangles of the recovered first-order and second-order correlation matrices and the true (simulated) first-order and second order correlation matrices. The averages are taken across several randomly generated synthetic datasets for each timeseries pattern (constant, random, ramping, and data; see *Synthetic data: simulating dynamic higher-order correlations*). The $x$-axes of each heatmap denote varying Laplace-shaped kernel widths (see Figs. S1 and S2), and the $y$-axes of each heatmap denote varying numbers of features ($K$) used in each of the synthetic datasets. A total of 10 synthetic datasets were generated for each number of features (row) and timeseries pattern (panel).

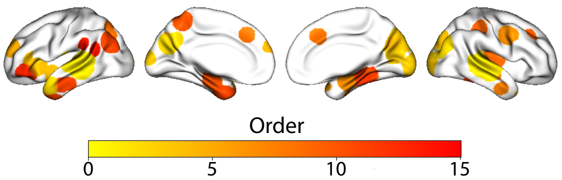


| Order 1 | Order 2 | Order 3 | Order 4 | Order 5 |
|---|---|---|---|---|
| voice | voice | cuneus | anterior insula | integrate |
| speech | precuneus | v1 | insula inferior | spatially |
| superior temporal | temporal | visual | inferior frontal | anterior hippocampus |
| listening | speaker | early visual | insula | occipital |
| temporal gyrus | temporal gyrus | primary visual | executive | recollection |
| sts | spoken | lingual gyrus | inferior | orbital |
| speech perception | heard | occiplital | frontal | hippocampal |
| spoken | sounds | lingual | pole | visual stream |
| superior | comprehension | visual cortex | frontal gyrus | visual cortex |
| temporal sulcus | speech perception | integrate | asd | visual |

| Order 6 | Order 7 | Order 8 | Order 9 | Order 10 |
|---|---|---|---|---|
| putamen | hippocampal | face recognition | monitoring | posterior insula |
| face ffa | hippocampus | recognition | error | insular |
| ffa | amygdala hippocampus | face | conflict | extrastriate |
| subsequent | anterior hippocampus | fusiform face | task | insular cortex |
| fusiform face | medial temporal | face ffa | frontal cortex | lateral occipital |
| images | parahippocampal | ffa | read | letter |
| selective | mtl | faces | errors | occipital |
| basal ganglia | temporal lobe | fusiform gyrus | insula anterior | visual |
| ganglia | parahippocampal cortex | hippocampal | color | painful |
| basal | lobe | medial temporal | demands | insula |

| Order 11 | Order 12 | Order 13 | Order 14 | Order 15 |
|---|---|---|---|---|
| memory encoding | angular gyrus | parahippocampal | inferior frontal | motor task |
| subsequent | angular | navigation | inferior | sensorimotor cortex |
| hippocampus | anterior temporal | objects | words | contralateral |
| hippocampal | temporal lobes | parahippocampal cortex | pseudowords | hand |
| associative | default | parahippocampal gyrus | frontal gyrus | sensorimotor |
| memory | default mode | anterior hippocampus | semantic | finger |
| lobe mtl | network dmn | encoding | reading | primary |
| mtl | self referential | memory encoding | chinese | spatially |
| retrieval | dmn | fusiform | word | tapping |
| episodic | default network | face ffa | orthographic | index finger |

**Figure S5: Top terms associated with the most strongly correlated nodes at each order, for the *intact* experimental condition.** Each color corresponds to one order of inter-subject functional correlations. The inflated brain plots display the locations of the endpoints of the 10 strongest (absolute value) correlations at each order, projected onto the cortical surface (Combrisson et al., 2019). The lists of terms display the top 10 Neurosynth terms (Rubin et al., 2017) decoded from the corresponding brain maps for each order. (Also see Fig. 6, top row, in the main text.)

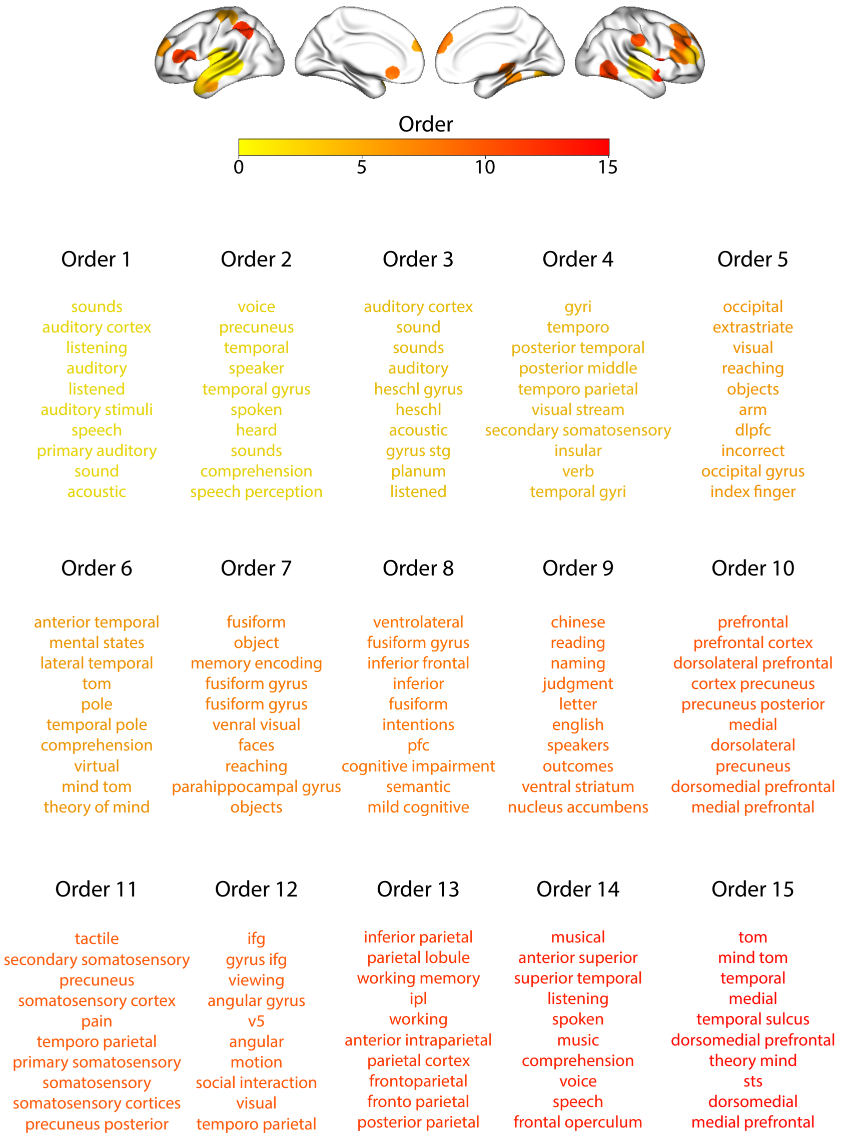


**Figure S6: Top terms associated with the most strongly correlated nodes at each order, for the *paragraph* experimental condition.** This figure is in the same format as Figure S5, but displays results for the paragraph-scrambled story listening condition. (Also see Fig. 6, second row, in the main text.)

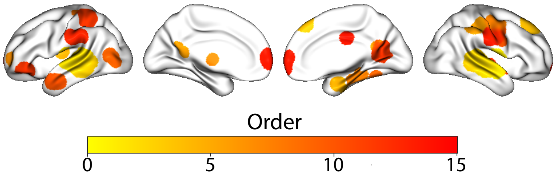


| Order 1 | Order 2 | Order 3 | Order 4 | Order 5 |
|---|---|---|---|---|
| listening | voice | temporal gyrus | anterior temporal | mild cognitive |
| speech | temporal gyrus | voice | expression | memory |
| voive | sounds | superior temporal | lateral temporal | medial temporal |
| sounds | listening | temporal sulcus | temporal lobes | lobe |
| speeck perception | speech | speaker | lobes | mtl |
| spoiken | spoken | sts | temporal lobe | retrieval |
| listened | audisotry | speech | fearful faces | episodic |
| audiory | superior temporal | listening | temporal | temporal lobe |
| superior temporal | listened | spoken | lobe | lobe mtl |
| temporal gyrus | speech perception | superior | neutral faces | episodic memory |

| Order 6 | Order 7 | Order 8 | Order 9 | Order 10 |
|---|---|---|---|---|
| inferior parietal | anterior temporal | parahippocampal | face | extrastriate |
| default mode | amygdala insula | episodic | fusiform face | occipitotemporal cortex |
| default | expression | parahippocampal gyrus | face ffa | occipitotemporal |
| inhibition | amygdala | hippocampus | ffa | videos |
| pcc | anterior prefrontal | hippocampal | fusiform | v5 |
| posterior cingulate | neutral faces | memory | faces | visual |
| response inhibition | temporal lobbe | medial temporal | fusiform gyrus | mt |
| medial | anxiety | anterior hippocampus | reaching | perception |
| autobiographical | fearful faces | episodic memory | occipital | fusiform |
| dmn | amygdala responses | temporal lobe | face recognition | objects |

| Order 11 | Order 12 | Order 13 | Order 14 | Order 15 |
|---|---|---|---|---|
| primary somatosensory | medial prefrontal | reaching | locations | medial |
| premotor | medial | retrosplenial | precuneus | posterior cingulate |
| anterior temporal | anterior temporal | visuomotor | body | pcc |
| somatosensory | prefrontal cortex | tools | supramarginal | medial prefrontal |
| premotor cortex | prefrontal | precuneus | videos | cortex precuneus |
| inferior parietal | semantic memory | videos | actions | referential |
| s1 | negative positive | touch | mt | self referential |
| ventral premotor | mpfc | cortex precuneus | video clips | cortex posterior |
| execution | autobiographical | navigation | video | autobiographical |
| somatosensory cortex | temporal lobe | retrosplenial cortex | occipitotemporal | hippocampus |

**Figure S7: Top terms associated with the most strongly correlated nodes at each order, for the *word* experimental condition.** This figure is in the same format as Figure S5, but displays results for the word-scrambled story listening condition. (Also see Fig. 6, third row, in the main text.)

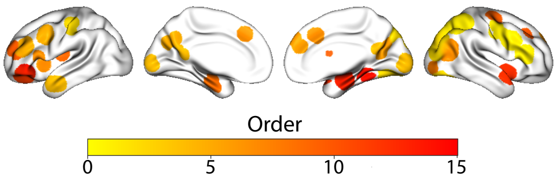


| Order 1 | Order 2 | Order 3 | Order 4 | Order 5 |
|---|---|---|---|---|
| cortex parietal | occipital | parietal | default | dorsolateral |
| intraparietal | visual | conflict | autobiographical | frontal |
| parietal cortex | etrastriate | task | default mode | middle frontal |
| ips | vision | visual cortex | posterior cingulate | motion |
| posterior parietal | occipital cortex | working memory | anterior temporal | mt |
| parietal | visual cortex | supplementary motor | dmn | verbal |
| intraparietal sulcus | visual stream | working | early visual | memory |
| symbbolic | motion | arithmetic | mental imagery | visual motion |
| superior parietal | v1 | supplementary | primary visual | dorsolateral prefrontal |
| prefrontal parietal | mt | anterior temporal | autobiographical memory | vision |

| Order 6 | Order 7 | Order 8 | Order 9 | Order 10 |
|---|---|---|---|---|
| locations | insual anterior | occipital | posterior insula | inferior occipital |
| navigation | error | precuneus | temporal lobe | anterior prefrontal |
| medial | anterior insula | letter | mtl | face |
| medial prefrontal | monitoring | inferior occipital | amygdala responses | selective |
| orienting | insula | visual | lobe mtl | face ffa |
| precuneus | conflict | cortex precuneus | insula | dlpfc |
| memory retrieval | frontal cortex | occipital cortex | lobe | ffa |
| location | insular | lateral occipital | mild cognitive | fusiform face |
| retrieval | taste | spatially | insular | dorsolateral |
| dorsomedial | errors | sulcus | cognitive impairment | cortex dlpfc |

| Order 11 | Order 12 | Order 13 | Order 14 | Order 15 |
|---|---|---|---|---|
| hippocampal | food | insula | amygdala hippocampus | s1 |
| hippocampus | ofc | taste | hippocampal | striatal |
| anterior hippocampus | cortex ofc | insular | hippocampus | outcomes |
| lobe | orbitofrontal cortex | insular cortex | anterior hippocampus | monetary |
| amygdala hippocampus | nucleus accumbens | unpleasant | medial temporal | memory encoding |
| parahippocampal gyrus | accumbens | pleasant | amygdala | accumbens |
| mtl | orbitofrontal | amygdala | parahippocampal | mesolimbic |
| medial temporal | value | amygdala insula | navigation | nucleus accumbens |
| temporal lobe | orbital | posterior insula | unpleasant | frontal |
| lobe mtl | hypothalamus | pain | episodic | money |

**Figure S8: Top terms associated with the most strongly correlated nodes at each order, for the *rest* experimental condition.** This figure is in the same format as Figure S5, but displays results for the resting state condition. (Also see Fig. 6, bottom row, in the main text.)

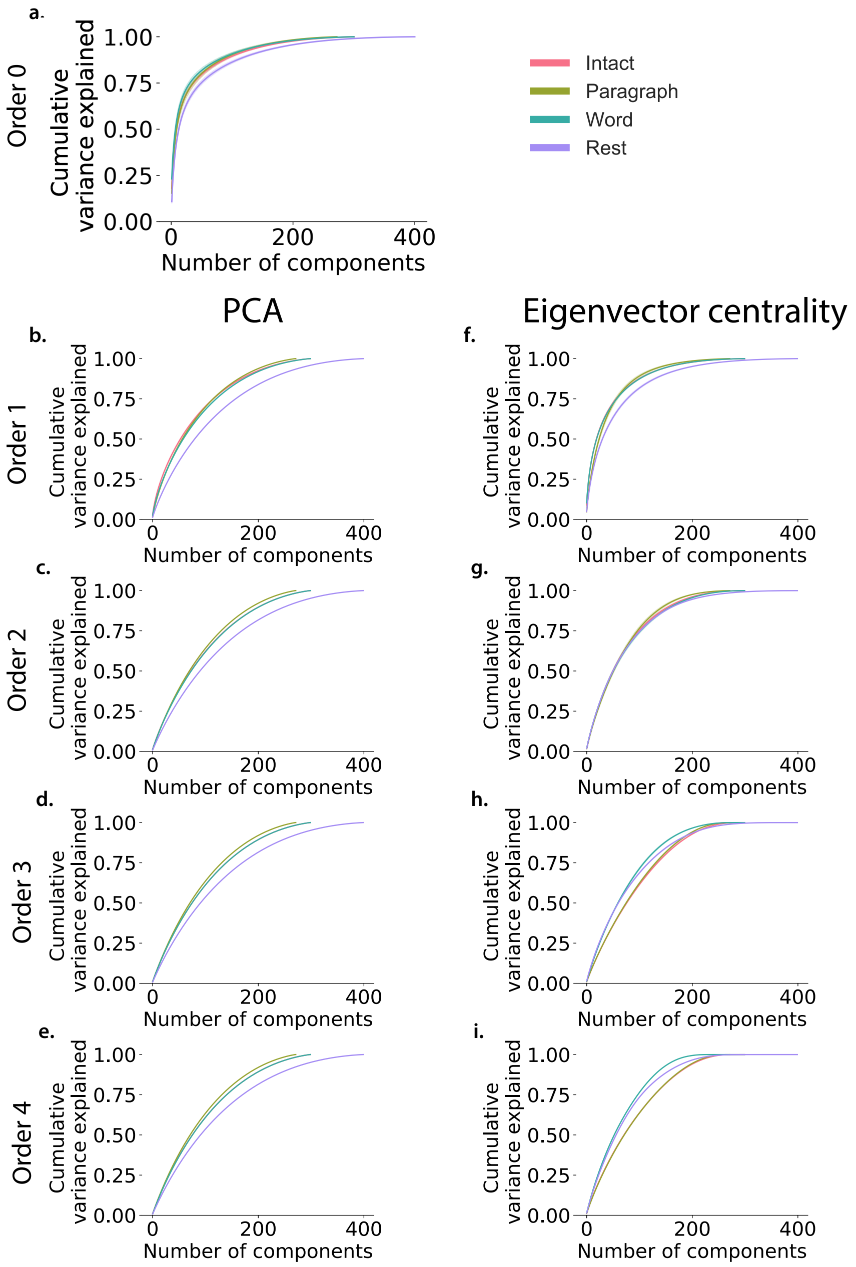


**Figure S9: Cumulative percent variance explained as a function of the number of principle components.** *Order* refers to the order of the dynamic correlations calculated. Principle components analysis was performed, and reduced independently for each subject. Maximum number of components varies with the total time for each condition (intact: 300; paragraph: 272; word: 300; rest: 400). **a. Cumulative percent variance as a function of number of components for Order 0.** PCA was performed on the raw activity patterns (Order 0). **b.-e. Cumlative percent variance as a function of number of components for Orders 1.-4.: PCA** Dynamic correlation were calculated for orders 1-4 using PCA to project each high-dimensional pattern of dynamic correlations onto a lower-dimensional space. **f.-i. Cumlative percent variance as a function of number of components for Orders 1.-4.: eigenvector centrality.** These panels are in the same format as Panel b.-e., but here eigenvector centrality has been used to project the high-dimensional patterns of dynamic correlations onto a lower-dimensional space.
